# Using the visual cliff and pole descent assays to detect binocular disruption in mice

**DOI:** 10.1101/2023.05.29.542767

**Authors:** Héctor De Jesús-Cortés, Teresa L.M. Cramer, Daniel A. Bowen, Francis Reilly-Andújar, Sophie Lu, Eric D. Gaier, Mark F. Bear

## Abstract

Amblyopia, a neurodevelopmental visual disorder characterized by impaired stereoacuity, is commonly modeled in animals using monocular deprivation (MD) during a critical period of visual development. Despite extensive research at the synaptic, cellular and circuit levels of analysis, reliable behavioral assays to study stereoscopic deficits in mice are limited. This study aimed to characterize the Visual Cliff Assay (VCA) and the Pole Descent Cliff Task (PDCT) in mice, and to evaluate their utility in detecting binocular dysfunction. Using these assays, we investigated the impact of clinically relevant manipulations of binocular vision, including monocular occlusion, pupillary dilation, and amblyopia induced by long-term MD. Our findings reveal that optimal performance in both the VCA and PDCT are dependent on balanced binocular input. However, deficits after MD in the VCA exhibited relatively small effect sizes (7-14%), requiring large sample sizes for statistical comparisons. In contrast, the PDCT demonstrated larger effect sizes (43-61%), allowing for reliable detection of binocular dysfunction with a smaller sample size. Both assays were validated using multiple monocular manipulations relevant to clinical paradigms, with the PDCT emerging as the preferred assay for detecting deficits in stereoscopic depth perception in mice. These findings provide a robust framework for using the VCA and PDCT in mechanistic and therapeutic studies in mice, offering insights into the neural mechanisms of binocular vision and potential interventions for amblyopia

## Introduction

Amblyopia (“lazy eye”) is a prevalent neurodevelopmental visual disorder characterized by impaired stereoacuity stemming from abnormal visual experience early in life (Bradfield, 2013; McConaghy and McGuirk, 2019). Monocular deprivation (MD) via unilateral eyelid suture during the critical period is the predominant paradigm to model and study amblyopia in animals (Wiesel and Hubel, 1963; Li et al., 2014). MD elicits an ocular dominance (OD) shift in V1, reflecting the loss of strength of the synaptic inputs from the deprived eye. Studies in cats and primates have shown that the primary locus of underlying functional plasticity is the synaptic neuropil of V1 itself. To gain an understanding of the molecular mechanisms for such plasticity, work increasingly has focused on the mouse. Limitations of the mouse as a model to understand primate vision are numerous and include limited binocular vision, poor spatial acuity, a poorly differentiated LGN with a high proportion of neurons with binocular receptive fields, and the absence of the columnar organization in V1 that is characteristic of carnivores and primates. Nevertheless, studies in mice have been instrumental in pinpointing changes at the molecular level that shift OD in V1 after MD. This work is directly relevant to the understanding of the most treatment-resistant form of amblyopia in humans, deprivation amblyopia, which arises from optical obstruction of the visual axis (e.g., by cataract) (Duffy et al., 2023).

Studies of MD in rodents have suggested novel treatment approaches to promote amblyopia recovery after the critical period. These include environmental enrichment (Sale et al., 2007), dark exposure (Montey and Quinlan, 2011), pharmacologic retinal silencing (Fong et al., 2016; Fong et al., 2021) and ketamine (Grieco et al., 2020; Venturino et al., 2021). Assays employed to assess treatment effectiveness typically include electrophysiological or optical measures of visual responses in V1, and behavioral estimates of monocular spatial acuity. Reliable behavioral assays to study stereoscopic deficits in mice, however, are needed.

To test stereoscopic depth perception of human infants, Gibson and Walk developed the visual cliff assay (VCA) (Gibson and Walk, 1960). The VCA is comprised of a glass plate atop a patterned floor with a sharp drop off. Subjects with functional stereoscopic vision are inherently averse to the “cliff” side and tend to spend more time on the “safe” side. The VCA has been adapted for use in several animal species to assess perception of stereoscopic depth (Cornwell et al., 1976; Kaufman, 1976; Morrison, 1982; Green et al., 1993; Witherington et al., 2005; Baroncelli et al., 2013; Rodkey, 2015). The assay has been used in rats to study the development of depth perception and the role of early-life visual experience (Kaufman, 1976; Morrison, 1982; Baroncelli et al., 2013; Rodkey, 2015), including the effect of MD (Baroncelli et al., 2013; Sansevero et al., 2019). Although it was initially questioned whether mice are capable of visually perceiving depth (Waugh, 1910), emerging evidence has demonstrated that mice are able to discriminate stereoscopic surfaces despite having only 40 degrees of binocular visual field (Drager, 1978; Heesy, 2004; Scholl et al., 2013; Samonds et al., 2019; Williams et al., 2021). Performance of mice in the VCA has been shown to impaired by restricting vision to one eye, albeit modestly (Mazziotti et al., 2017).

Recently, a modified version of the VCA has been described, called the pole descent cliff task (PDCT) (Boone et al., 2021). In this task, mice descend a vertical pole to a conical platform which engages the upper visual field, where there is more binocular overlap in visual representations. The vertical pole is suspended over a glass plate divided into four quadrants; three of the four quadrants appear as a visual cliff using stereoscopic depth cues. Neurotypical mice, viewing with both eyes, will exit the pole onto the “safe” quadrant. Acutely closing one eye results in errors that increase as the apparent depth of the cliff quadrants increases. Although promising, performance has not yet been shown to be sensitive to disruption of stereopsis caused by deprivation amblyopia following long term MD (LTMD) in mice.

Robust and reliable behavioral assays of binocular vision are clearly needed to validate functional outcomes observed in experimental treatment paradigms in mice. We therefore set out in the current study to investigate the utility of the VCA and PDCT to probe deficits in binocular vision in mice caused by long-term MD.

## Materials and methods

### Animals

All experimental procedures followed the guidelines of the National Institutes of Health and the Association for Assessment and Accreditation of Laboratory Animal Care International and adhered to protocols approved by the Committee on Animal Care at MIT. Female and male C57BL/6 mice were housed in groups of 2-5 same-sex littermates beginning at P21 after being bred in the MIT animal colony. Mice had access to food and water *ad libitum* and were kept in a temperature-controlled room and maintained on a 12-hour light/dark cycle. Animals with any apparent congenital ocular abnormalities were excluded.

### Eyelid Suture

For monocular deprivation experiments, eyelid suture (or sham eyelid suture) occurred at postnatal day (P) 21. Mice were anesthetized via inhaled isoflurane (1-3% in oxygen). Both eyes were inspected for abnormalities and kept lubricated using single-use artificial tears. The right eyelid was closed with two horizontal mattress sutures using 7-0 Prolene (Ethicon, polypropylene, 8648 G). Toenails on both forepaws and hind paws were trimmed. Littermate controls designated as “sham” had their eyes reopened immediately after being sutured shut. For reopening of the eye following long-term MD, mice were anesthetized and the sutures removed from the deprived eye. Sham controls were kept under isoflurane for an equivalent amount of time. Mice whose eyelids were not fully shut at the time of eye reopening or demonstrated any apparent ocular injury were excluded.

### Visual Cliff Assay

The VCA was conducted in a quiet, temperature-controlled room during the 12-hour light cycle. All mice had their whiskers trimmed 1 hour before performing the VCA to minimize use of non-visual cues during the test. The apparatus was modeled after that published by Han et al. (2017) and consisted of an open field behavioral box with transparent plexiglass walls and floor (42cm x 42cm x 30.5cm), positioned on the edge of a laboratory bench with half the box on the table top (the “safe” side), and the other half extended over a floor 80 cm below (the “cliff” side) (Han et al., 2017). A black and white striped cover was placed on the bench top and dropped down vertically to the floor to create the illusion of a cliff. At the start of each trial, animals were placed on a petri dish in the middle of the box facing the safe side, as illustrated in **Figure 1A**. A 1080P wide angle ELP 100fps USB camera with infrared capability was kept 52cm above the plexiglass floor of the box and connected to a computer to record each session. The mice were brought to the same room as the apparatus and habituated for at least 30 minutes in their home cage before beginning each trial. The trial began after the animal stepped off the petri dish and ended 2-5 minutes later. The surface of the plexiglass chamber was cleaned with peroxigard between all trials. In all test-retest experiments, the chamber was rotated between trials for each animal to mitigate contributions by visual cues not related to the cliff.

**Figure 1.**
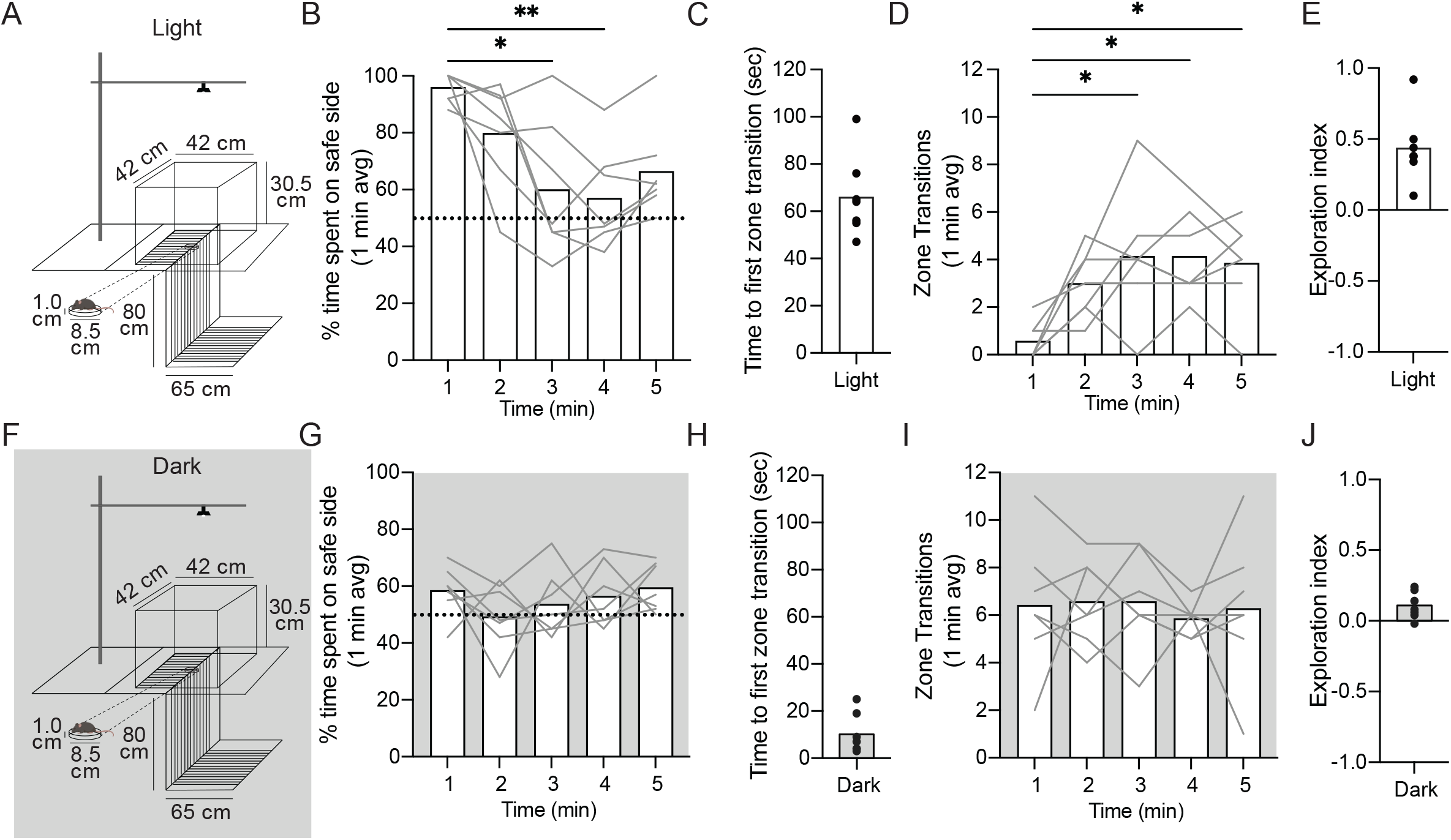
VCA performance requires visual input. **A**. Graphical design of the VCA with measurements of the box and table that were used. Animals were placed on a petri dish (2cm height) in the middle of the platform facing the safe side and allowed to explore the arena for 5 minutes. **B**. Percent time spent on safe side (per minute) plotted as a function of test minute. Dotted line (y=50%) represents chance performance. **C**. Time to first zone transition from safe to cliff side plotted as a function of test minute. **D**. The total number of transitions between zones plotted as a function of test minute. **E**. Exploration index ([time on safe – time on cliff]/ total time) over the 5-minutes test. **F**. Graphical design of the VCA in the dark using a different cohort of animals. **G**. Percent time spent on safe side in the dark (per minute) plotted as a function of test minute. Dotted line (y=50%) represents chance performance. **H**. Time to first zone transition from safe to cliff side in the dark plotted as a function of test minute. **I**. The total number of transitions between zones in the dark plotted as a function of test minute. **J**. Exploration index over the 5-minutes test in the dark. Mean values are represented as bars for each test minute in each condition. Individual animals are represented as grey lines. Dunn’s multiple comparison representations of statistical significance compared to the first minute in each condition: **\*** <.05, **\*\*** <.01.

Visual cliff performance was scored manually using a stopwatch to calculate the percent time the mouse spent on the safe side, the number of zone transitions between sides, and the time of the first transition to the cliff side by an experimenter blinded to condition. A zone transition was defined as all four limbs of the mouse crossing to the next zone. Manually scored metrics were confirmed using the open-source pose estimation software ezTrack (https://eztrack.studio), which was also used to measure total distance traveled and locomotion for each animal.

### Pole Descent Cliff Task

All PDCT experiments were conducted in a quiet, temperature-controlled room during the 12-hour light cycle. The apparatus was modeled after Boone et al. (2021) and consisted of an open field behavioral box (42cm x 42cm x 30.5cm) placed above museum glass. A round wooden dowel (pole) 3.5 cm in diameter was screwed into a 75ml Erlenmeyer Glass Flask 9 cm in diameter at the bottom (total combined height of 52 cm). The pole and flask were covered with anti-slip silicone sleeves, texturized with a shelf and cabinet grip liner and spray painted white. The flask was weighted with sand and a spacer was glued to the center bottom of the flask, which allowed it to hover 1.5 cm above the museum glass. A platform covered in a 2.5 × 2.5 cm sized black and white checkerboard pattern was placed 23 cm below the glass. Another platform, covering one quadrant of the glass, was covered in checkerboard pattern and placed immediately below the glass. The pole was positioned in the center of the museum glass. A 1080P wide angle ELP 100fps USB camera with infrared capability was kept 65cm above the plexiglass floor of the box and was connected to a computer to record each session.

Prior to testing day, mice were individually handed for a minimum of 2 days, 5 min each. On the day of testing, whiskers were removed, and mice were brought to the behavioral testing room for a minimum of 30 min habitation in their home cage. Before beginning each trial, mice were allowed to freely explore the open field behavioral box for 5 min. During this habitation, the checkerboard patterned platform was placed directly right below the museum glass, and the pole was positioned in the center of the behavioral box. At the end of each habituation session, each mouse was allowed to descent the pole once. For the trial, the platform was placed 23 cm below the glass, and the position of the nearest quadrant was rotated every 1-2 mice to eliminate any side preference due to external cues. Mice were placed on the top of the pole facing downwards to motivate descent and videos were acquired of all behavior for later analysis. Between running of each animal, the chamber and pole were cleaned with peroxigard to eliminate odor cues. Cohorts of male mice tested to completion, then cohorts of female mice were tested thereafter.

For video analyses, manual scoring of the quadrant (safe/unsafe) mice exited to, and time taken to exit the pole (descent time) was performed. Experimentation and video analysis is done experimenter-blind to group/intervention.

### Experimental design and statistical tests

Pilot experiments were run for each study using 3-5 animals (not included in final data), and necessary sample sizes were calculated using the G* power (Faul et al., 2007). All other statistical testing was performed using GraphPad (Prism version 9.0 for Mac, GraphPad Software, San Diego, California USA). Normality of all datasets and groups were tested using the normality test Shapiro-Wilk. Datasets containing non-normally distributed data were tested using non-parametric tests (Kruskal-Wallis, Friedman or Wilcoxon tests). Normally distributed data were analyzed by ANOVA or *t* tests. *p*<.05, adjusted for multiple comparisons, was considered statistically significant in all cases.

## Results

### The contribution of vision to the VCA

To characterize the VCA, we first quantified the inherent tendency of mice to stay on the “safe” side under normal lighting conditions. Female and male mice (P45-58, n=7) were placed on the middle of the apparatus (**Figure 1A**) facing the “safe” side and allowed to freely explore the arena for 5 minutes as previously described (Han et al., 2017; Mazziotti et al., 2017).

Sessions were video recorded from above and behaviors were scored by percent time spent on the safe side by each minute (**Figure 1B**), the time to first zone transition (**Figure 1C**), and the number transitions between zones by minute (**Figure 1D**). On average, animals spent 96.0±4.8% (95% CI) of the first minute on the safe side. A non-parametric repeated Friedman test showed a main effect of test minute on the proportion of time spent on the safe side over the 5-minute test (*p* =.0060). Pairwise comparisons revealed that time spent on the safe side decreased significantly to 60.0±22.2% (Δ=37.5%; *p*=.0162) and 57.0±22.2% (Δ=40.6%; *p*=.0021) during the third and fourth minutes, respectively (**Figure 1B**). On average, animals made the first zone transition to the cliff side at 66.0±15.9 seconds (**Figure 1C**). Repeated Friedman test showed a main effect of test minute on the number of zone transitions (*p*=.0088). We observed a significant increase in the number of zone transitions by the third (1^st^ min: 0.6 ± 0.7, 3^rd^ min: 4.1 ± 2.5; *p*=.0211), fourth (4.1±1.7; *p*=.0162), and fifth minutes (3.9±1.8; *p*=.0211) (Fig. 1D). The exploration index (EI; difference between time on safe and cliff sides divided by the total time), a standard single measure of VCA performance, was 0.44±0.23 (**Figure 1E**).

To evaluate the contribution of vision, we tested VCA performance in complete darkness in a separate cohort of animals (P45-58, n=7) (**Figure 1F**). On average, animals spent 58.6±8.2% of the first minute on the safe side (**Figure 1G**), significantly less than in the light (*p*=.0006; Mann Whitney test). Repeated measures ANOVA showed no effect of test minute on the proportion of time spent on the safe side (*p*=.3765) (**Figure 1G**), that ranged between 49.3– 59.6% over the 5-minute test. On average, animals made the first zone transition to the cliff side at 10.1±7.9 seconds (**Figure 1H**), significantly sooner than in the light (*p*<.0001, unpaired *t* test). Repeated measures ANOVA showed no effect of test minute on the number of zone transitions (*p*=.8995) (**Figure 1I**). The EI was significantly decreased in the dark (0.11±0.09) compared to the light (*p*=.0006, unpaired *t* test) (**Figure 1E vs J**). Taken together, these data unambiguously indicate that vision is required for normal performance on the VCA in mice, and that visual cues deter zone transitions in the first 2 minutes of the test.

### Test-retest stability of VCA performance

Recognizing that experiments involving a treatment or intervention may benefit from multiple runs in the VCA in the same animals, we next examined test-retest reliability. We modified the assay to only test animals for two minutes to avoid habituation to the cliff side in the first trial (see **Figure 1**). A new cohort of female and male mice (P48-68, n=20) was run 4 times at 1-week intervals. Repeated measures Friedman test showed a borderline, but ultimately statistically non-significant effect of trial number on percent of time spent on the safe side (*p*=.0589) (**Figure 2A**). Given the marginal effect of trial number and our objective to assess repeatability over weeks, we examined pairwise comparisons. Between week 0 and week 1, there was no significant change in time spent on the safe side (*p*=.0825). There was a main effect of trial number on the time to first zone transition (*p*=.0248) that was carried by a decrease between the first and second trial (**Figure 2B**, *p*=.0212). There was a main effect of trial number and the number of zone transitions (*p*=.0218) with no pairwise differences with the first trial (**Figure 2C**, *p* values>.1501). Thus, VCA performance remains relatively stable over weeks.

**Figure 2.**
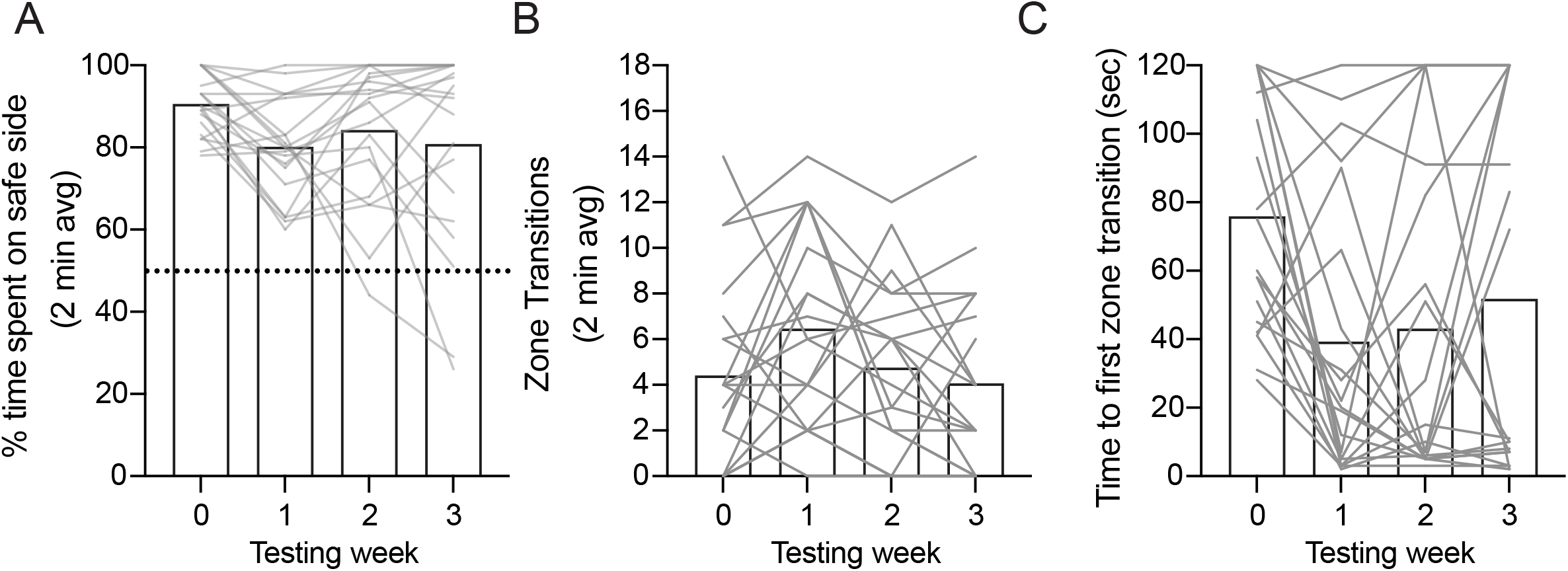
Test-retest stability of VCA performance. **A**. Percent time spent on safe side (average over each 2-minute trial) plotted as a function of testing week (1-week trial interval). Dotted line (y=50%) represents chance performance. **B**. Time to first zone transition from safe to cliff side plotted as a function of testing week. **C**. The total number of transitions between zones plotted as a function of testing week. Mean values are represented as bars for each test week. Individual animals are represented as grey lines. Dunn’s multiple comparison representations of statistical significance compared to week 0: **\*** <.05.

### Acute disruptions of binocularity impair VCA performance

Knowing that VCA performance requires vision, we next tested the contribution of binocularity in the 2-minute test. VCA performance has previously been shown to drop under monocular conditions (with one eye sutured closed) in both rats (Baroncelli et al., 2013) and mice (Mazziotti et al., 2017), so we began by attempting to replicate this result. Male and female mice (P42-60, n=11 per group) had one or both eyes sutured closed under isoflurane anesthesia 1-3 hours prior to VCA testing. Control (sham) animals underwent the same manipulations, but the eyes were immediately re-opened. Eye closure decreased performance in a dose-dependent manner across all metrics (**Figure 3A–D**). ANOVA showed a main effect of eye closure on percent time spent on the safe side (*p*<.0001). Time spent on the safe side decreased by 14.0% with one eye closed (*p*=0.0082) and 29.7% with both eyes closed (*p*<.0001) (**Figure 3A**). There was a main effect of eye closure on time to first zone transition (*p*=.0013) with a decrease of 38.5±8.8 seconds with one eye closed (*p*=.0068) and 11.2±6.7 seconds with both eyes closed (*p*=.0025) (**Figure 3B**). There was a main effect of eye closure on the number of zone transition (*p*=.0006) (**Figure 3C**) with an increase of 5.3±2.5 with one eye closed (*p*=.0085) and 6.4±2.1 with both eyes closed (*p*=.0019). The dose-dependence of eye closure on VCA performance corroborates the dependence of the VCA on vision (**Figure 1**), and also suggests a contribution of binocular function. Considering that differences in movement may confound VCA performance comparisons between groups, we used open-source pose estimation software (EzTrack (Pennington et al., 2021) to quantify the total distance traveled during the 2-minute test. There was a main effect of eye closure on the total distance traveled (*p*=.0006) (**Figure 3D**). Pairwise comparisons revealed a significant increase of 501 cm (45%) in total distance traveled when both eyes were closed (*p*=.0009), but no significant change when only one eye was closed (248 cm; *p*=.0990). Therefore, we can at least partially attribute VCA performance deficits under monocular conditions to a lack of binocularity.

**Figure 3.**
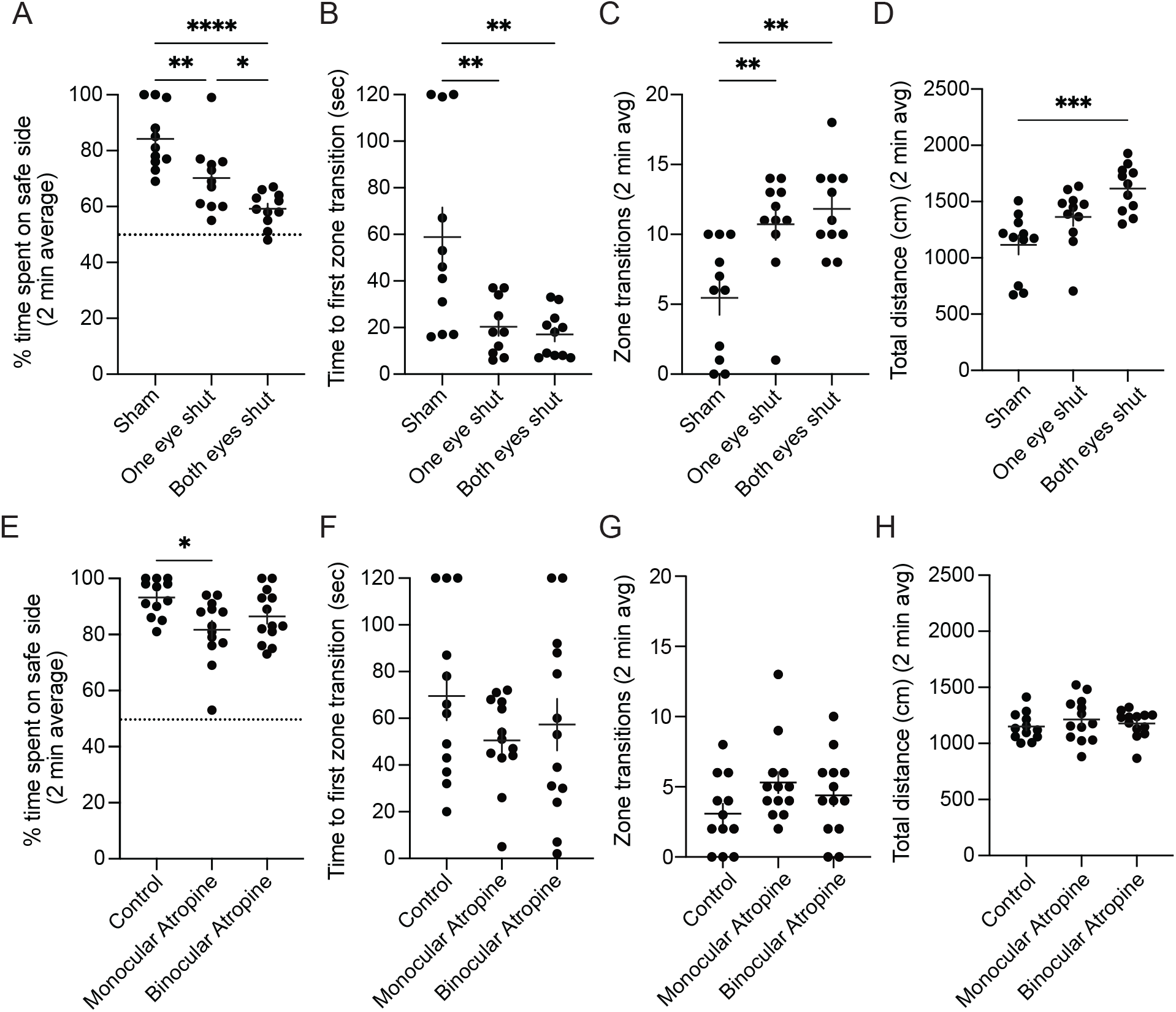
Disruptions of binocularity impair VCA performance. (**A–D**) Animals performed the 2-minute VCA with neither (sham), one or both eyes sutured shut. **A**. Percent time spent on safe side plotted for each eye closure group. Dotted line (y=50%) represents chance performance. **B**. Time to first zone transition from safe to cliff side plotted for each eye closure group. **C**. The total number of transitions between zones plotted for each eye closure group. **D**. Total distance traveled during the test plotted for each eye closure group. (**E–H**) Animals performed the 2-minute VCA after neither, unilateral or bilateral pupillary dilation via atropine ointment administration. **E**. Percent time spent on safe side plotted for each atropine group. Dotted line (y=50%) represents chance performance. **F**. Time to first zone transition from safe to cliff side plotted for each eye atropine group. **G**. The total number of transitions between zones plotted for each atropine group. **H**. Total distance traveled during the test plotted for each atropine group. Individual animals are represented as circles and mean ±SEM as black horizontal and vertical lines. Dunn’s multiple comparison representations of statistical significance between groups: **\*** <.05, **\*\*** <.01, **\*\*\*** <.001, **\*\*\*\*** <.0001.

Monocular occlusion significantly narrows the visual field particularly in mice and could influence VCA performance, which is globally dependent on vision (**Figure 1**). To better delineate whether VCA performance is dependent on balanced binocular inputs rather than a narrowing of the visual field, we next tested the influence of optical manipulations. Spherical aberration degrades spatial acuity and stereopsis, particularly with interocular disparity (Jiménez et al., 2008). The optics of the murine eye impart significant susceptibility to optical aberration as a function of pupil size (Gomes et al., 2023), likely owing to the relative sphericity of the lens. We applied atropine to dilate the pupil of one or both eyes and tested animals for 2 minutes in the VCA ≥60 minutes later (P38-42, n=12–13 per group, **Fig. 3E–H**). ANOVA showed a main effect of atropine on time spent on the safe side (*p*=.0160). Pairwise comparisons revealed that monocular atropine decreased the percent time spent on the safe side by 11.5% (*p*=.0083) whereas binocular atropine imparted no significant change (*p*=.1464) (**Figure 3E**). Atropine imparted no main effect on the time to first zone transition (*p*=.3551) (**Figure 3F**) or the number of zone transitions (*p*=.1826) (**Figure 3G**). In contrast to eye closure, atropine imparted no change in total distance traveled during the test (*p*=.5630) (**Figure 3H**). Therefore, the effect of atropine on time spent on the safe side can be attributed to binocular imbalance. These data demonstrate that disruption of balanced binocular input is sufficient to impair VCA performance.

### Amblyopic mice demonstrate binocular dysfunction in the VCA

We next tested the effects of amblyogenic rearing. Female and male mice underwent 3 weeks of long-term MD (LTMD) or sham eyelid closure/opening from P21–P42 and were tested in the VCA 1-3 hours after eye opening (**Figure 4A**). To ensure the study was adequately powered to detect meaningful group differences, an interim power analysis was performed after testing the first cohort of Sham control (n= 14) and LTMD (n= 14) animals. Based on this analysis, with a Type I error rate set at 5% (two-sided) and a desired power of 80%, a sample size of 44 animals per group was determined. Amblyopic mice (Sham control n=44, LTMD n=52) showed impaired performance across all VCA metrics (**Figure 4B–D**). LTMD decreased time spent on the safe side by 7.5% compared to control animals (*p*=.0070) (**Figure 4B**). LTMD decreased the time to first zone transition by 16.5 seconds (20.6%) (*p*=.0243) (**Figure 4C**) and increased the number of zone transitions (*p*=.0393) (**Figure 4D**). LTMD had no effect on total distance traveled (*p*=.6218) (**Figure 4E**), suggesting that amblyopia does not increase ambulatory movement as occlusion does. Together, these data demonstrate that LTMD imparts an amblyopic deficit in binocular function in mice to which multiple VCA metrics are sensitive.

**Figure 4.**
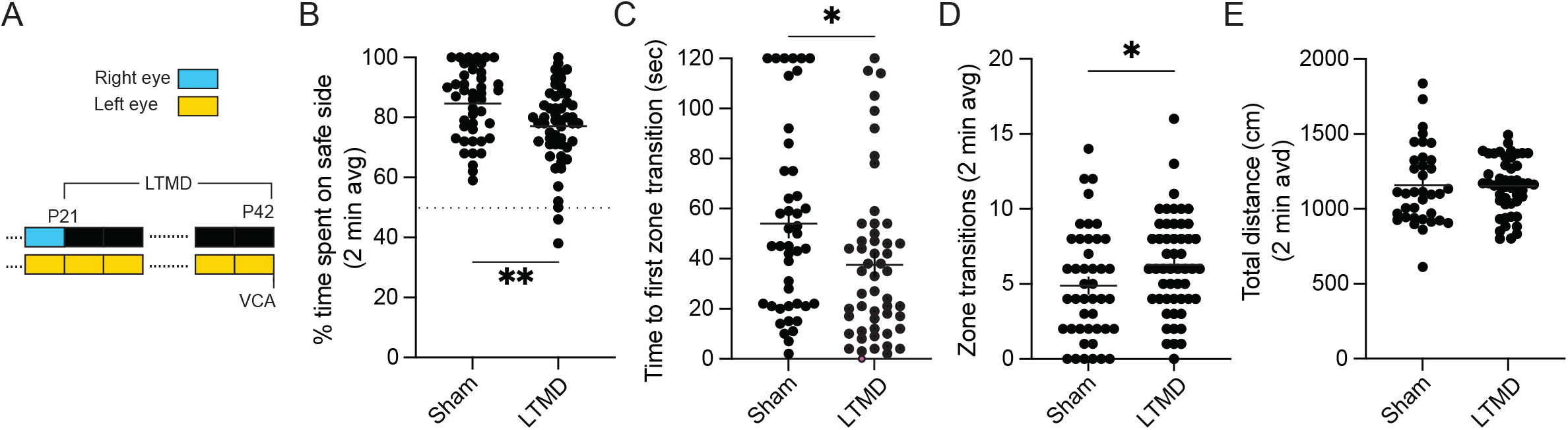
LTMD causes stable binocular dysfunction in the VCA. **A**. Experimental timeline showing LTMD (or sham closure) performed from P21-P42. VCA was performed 1-3hrs after eye opening (**B–E**). **B**. Percent time spent on safe side plotted for each group. Dotted line (y=50%) represents chance performance. **C**. Time to first zone transition from safe to cliff side plotted for each group. **D**. The total number of transitions between zones plotted for each group. **E**. Total distance traveled during the test plotted for each group. Individual animals are represented as circles and mean ±SEM as black horizontal and vertical lines. Comparison representations of statistical significance between groups (Mann-Whitney) or between tests (paired Wilcoxon): **\*** <.05, **\*\*** <.01.

### The PDCT reveals visual deficits following acute disruption of binocularity

Previous behavioral work has demonstrated that mice use binocular information in the PDCT to infer depth and exit the pole onto the nearest quadrant (Boone et al., 2021). We attempted to replicate these results and began by quantifying the overall tendency of mice to exit the pole. Female and male mice (P75-P110, n=9) were placed on the pole facing downwards to motivate descent (**Figure 5A**). Sessions were video recorded from above and behaviors were quantified as percent exits. Across 10 interleaved trials each lasting 3 minutes, the percentage of mice that exited the pole onto any of the four quadrants dropped significantly (**Figure 5B**). While 100% of mice exited the pole on the first trial, only 33.3±16.7% exited the pole on the fifth trial and no mice exited the pole on the tenth trial. These findings differ from published results obtained using the PDCT, where the fractional success of mice exiting the pole to the nearest quadrant was determined from 10–15 interleaved trials (Boone et al., 2021). We thus modified the assay to test animals in a single trial only.

**Figure 5.**
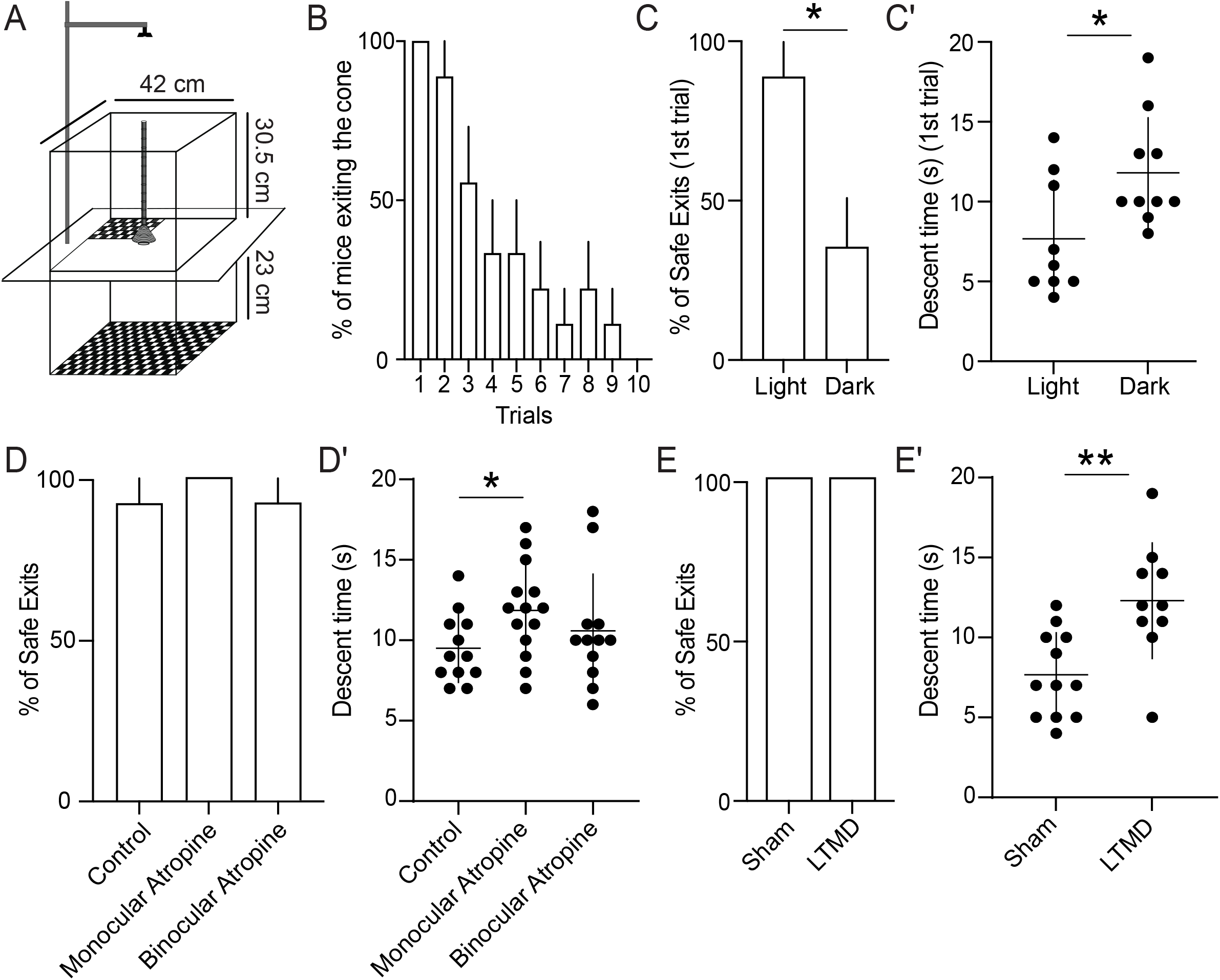
Disruptions of binocularity and LTMD impair descent time in PDCT. **A**. Graphical design of the PDCT apparatus with measurements that were used. Mice descend a pole positioned above a cone (9 cm diam) placed on a glass plate. One quadrant is positioned 1.5 cm below the cone, while the remaining three are 24.5 cm below the cone. All quadrants are covered with black and white checkerboard. **B**. % of mice that exit cone onto any quadrant across 10 interleaved trials each lasting 3 minutes. **C**. % of safe exits (exit to quadrant 1.5 cm below cone) under normal lighting conditions and in complete darkness. **C’**. Descent time (the time taken to descend the pole) under normal lighting conditions and in complete darkness. **D**. % of safe exits of animals after neither, unilateral or bilateral pupillary dilation via atropine ointment administration. **D’**. Descent time plotted for each atropine group. **E**. % of safe exits of sham and LTMD animals. **E’**. Descent time plotted for sham and LTMD animals. Bar graphs: mean ± SEM, Scatter plots: mean ± SD, and individual animals are represented as circles. Comparison representations of statistical significance between groups (unpaired t-test): **\*** <.05.

To evaluate the contribution of vision, we next tested PDCT performance under normal lighting conditions and in complete darkness using two separate cohorts of animals (P65-P95, n=9-10 per group) (**Figure 5C**). On average, 88.9±11.1% of animals exited the pole onto the nearest quadrant under normal lighting conditions, while only 36.3±16.1% of animals exited onto the nearest quadrant in darkness (*p*=0.015, unpaired t-test). These findings indicate that vision is required for performance in the PDCT. While observing animal behavior over the course of the assay, we noticed that animals tested in darkness took longer to descend the pole. The lack of visual cues in darkness prevents mice from using vision to gauge distances, which likely results in slower descent times. We quantified the descent time and found that animals tested in darkness took significantly longer to exit the pole (11.8±3.6 seconds) than animals tested in the light (7.6±3.7 seconds, *p*=0.021, unpaired *t* test). Taken together, these findings demonstrate that vision is required for descending and exiting the pole onto the nearest quadrant in the PDCT.

We next tested whether binocularity is required for PDCT performance. Using the VCA, we had found that monocular atropine and eye suturing produce comparable deficits in performance, as measured by time spent on the safe side (**Figure 3**). Because atropine is simple to apply, does not narrow the visual field, and may better approximate the effect of amblyopia, we focused on this manipulation in the PDCT. Atropine was applied to one or both eyes and the animals were tested in the PDCT ∼60 minutes later when pupil dilation as maximal (P55-65, n=13–14 per group, **Figure 5D-D’**). Surprisingly, atropine imparted no main effect on mice exiting the pole onto the nearest quadrant (control: 92.3 ± 0.7%, monocular atropine 100%; binocular atropine 92.8 ± 0.7%, *p*=0.60, **Figure 5D**). However, pairwise comparisons revealed that monocular atropine significantly increased the descent time compared to controls (*p*=0.02, unpaired *t* test), while binocular atropine did not (*p*=0.38, unpaired *t* test, **Figure 5D’**). These findings indicate that the PDCT metric “safe/unsafe exit” may not be sensitive enough to detect subtle imbalances in binocular vision. In contrast, mild disruption of balanced binocular input is sufficient to impair pole descent time in the PDCT.

### Amblyopic mice demonstrate binocular dysfunction in the PDCT

We next tested the effects of amblyogenic rearing in the PDCT. Female and male mice (n=8-10 per group) underwent 3 weeks of LTMD or sham eyelid closure/opening from P21–P42 and were tested in the VCA 1-3 hours after eye opening as shown in **Figure 4A**. We found that LTMD imparted no effect on mice exiting the pole onto the nearest quadrant (Sham: 100%, LTMD: 100%, **Figure 5E**). However, pairwise comparison revealed that LTMD significantly increased the descent time compared to sham controls (*p*=0.008, unpaired *t* test, **Figure 5E’**).

These findings indicate that LTMD is sufficient to impair pole descent time in the PDCT.

## Discussion

We characterized the VCA and PDCT as methods to study binocular perceptual deficits in mice. Mice are the preferred species to study experience-dependent neuroplasticity of visual processing despite important differences from humans with respect to neuroanatomy and circuitry of the visual system (Hooks and Chen, 2020). Mechanistic studies in mice are best validated through behavioral studies that demonstrate functional relevance. Here, we provide evidence that the VCA and PDCT rely on balanced binocular inputs for optimal performance in mice. Both assays have advantageous features including relatively fast execution, no need for operant training, and readout through several simple metrics. However, the assays also have important limitations, as evidenced through our results, that should be weighed carefully when considering their use.

### Impairments in binocular function are detected by the VCA, but effect sizes are modest

Monocular occlusion, pupillary dilation (and consequential optical aberration), and amblyopia all degraded performance in the VCA, supporting the role of binocular sensory integration as required for the task. Monocular pupillary dilation significantly reduced % time spent on the safe side whereas bilateral atropine did not (**Fig. 3E**) highlighting the need for balanced visual input in binocular integration. Monocular blur more significantly degrades stereopsis as compared with binocular blur (Westheimer and McKee, 1980; Legge and Gu, 1989). However, the manipulations that disrupted binocular function did not drive VCA performance to levels we observed running animals in the dark (**Fig. 1G–J**) or with both eyes closed (**Fig. 3A–C**). Manipulations of binocular function in our studies, such as unilateral pupillary dilation and amblyopia after MD, may only partially disrupt binocular function.

However, this is unlikely to be the sole explanation for the difference, as the unambiguous loss of binocular function caused by closing one eye similarly did not reduce performance to the levels observed in darkness or with both eyes closed. Another mode by which animals can detect depth is by discerning changes in monocular cues with movement, called motion parallax (Ellard et al., 1984; Ellard et al., 1986). Motion parallax may account for the partial effects of complete disruption of binocularity on VCA performance and is an unavoidable potential confounder in the VCA.

The main limitation for use of the VCA to detect deficits in binocular function in mice is the relatively small effect sizes (7-14%), which we found consistently across experiments and VCA metrics. Our observations are consistent with those reported previously in mice (Mazziotti et al., 2017), although that study reported a larger effect of MD (∼20%) when tested in younger animals during the critical period. Modest effect sizes limit the utility of this assay to detect partial improvement after an experimental treatment. Based on our observations in amblyopia caused by 3-week LTMD from P21 (**Fig. 4**), to achieve 80% power, hypothetical treatments restoring 100%, 75%, 50% binocular function would require 42, 73, and 165 animals per group to statistically demonstrate an effect. We made multiple iterations of and adjustments to the VCA to address this limitation without success.

### The PDCT is a novel approach to study deficits in binocular vision in amblyopic mice

While the VCA has a rich history of use in species from rodents to humans (Gibson and Walk, 1960; Fox, 1965; Baroncelli et al., 2013), only one prior study has described and applied the PDCT to evaluate binocular vision in depth perception in mice (Boone et al., 2021). To our knowledge, we are the first to attempt to replicate their findings and use the assay to study the effect of amblyogenic rearing. Under normal lighting conditions, we found that animals preferentially exited the pole onto the “safe” quadrant, while animals tested in darkness did not (**Fig 5C**). In agreement with Boone et al., we thus find that vision is required for targeting the nearest platform in the PDCT. We additionally found that animals tested in darkness require longer times to descend to the pole (**Fig 5D**). Of note, Boone et al. did not test animals in darkness. The absence of light likely makes the descent more challenging and slower by preventing mice from using vision to gauge distances, detect obstacles, and maintain balance. Additionally, without visual input, mice may rely more on other senses, such as touch, which could be less effective for quick navigation compared to visual guidance in a well-lit environment. Our observed increase in descent time is thus not surprising.

To test whether binocular vision is required for PDCT, we utilized pupillary dilation and amblyogenic rearing. Both paradigms degraded descent time performance in the PDCT but did not affect the tendency of mice to exit onto the nearest platform (**Fig 5D-E**). Thus, while we found the tendency of mice to exit onto the nearest quadrant to be dependent on vision, we found only descent time to be dependent on balanced binocular vision. These findings differ from published results obtained using the PDCT (Boone et al., 2021), where lack of binocularity caused by monocular occlusion impaired identification of the nearest platform without affecting descent time. These differences in behaviors may be attributed to how each paradigm affects visual processing and depth perception. Monocular occlusion eliminates binocular vision and substantially restricts the visual field. This absence of binocular function hinders the mice’s ability to accurately judge distances between platforms and determine the closest quadrant.

Despite this, mice with monocular vision can unambiguously detect contrast, light, and motion with normal acuity. As a result, monocular cues alone may provide sufficient information for spatial navigation, which explains why descent time was unaffected in the Boone et al. (2021) study. Unlike occlusion, monocular pupillary dilation introduces optical aberration in one eye, bringing the inputs from the two eyes into rivalry or conflict (which is resolved when both eyes receive atropine). Rivalry and visual confusion do not completely eliminate binocular fusion required for depth perception but rather diminish it. Similarly, and even though amblyopia following LTMD drastically affects the development of normal binocular vision, visual input from the deprived eye is not completely eliminated as it is following monocular occlusion. The remaining visual input is blurred or compromised, diminishing binocular fusion. At high binocular disparities found close to the platform, the remaining binocular function is likely sufficient to help mice accurately identify and target the nearest platform. At lower binocular disparities found near the top of the pole, depth perception relies heavily on subtle differences between the images seen by each eye. Here, the overall diminution of binocular function may result in slower descent times as mice navigate with reduced confidence in their depth perception. The discrepancies between our findings and those of Boone et al. are thus likely due to these methodological differences.

While we could not detect subtle imbalances in binocular vision caused by atropine application or LTMD using the metric “safe/unsafe exit”, we did find pole descent time to be sensitive enough to detect these subtle imbalances. As LTMD in mice has been the standard model to study the pathophysiology and novel treatment approaches of amblyopia, our findings strongly support the use of descent time in the PDCT to study stereoscopic deficits in mice. The major advantage of using descent time in the PDCT is its relatively large effect size (43-61%), which we found when utilizing atropine or LTMD to disrupt binocular vision (**Fig 5E**). This necessitated the use of <15 animals to make reliable comparisons, even when binocular vision is only partially disrupted.

Strengths of our study include the systematic approach to characterize VCA performance, replicate the PDCT, and determine their dependencies on binocular vision in mice. Our primary experiments are well-powered and thus provide a guide for future experiments to employ these assays. The pattern of behavior was consistent across experiments in our study. Manipulations that reduced binocular function resulted in decreased time spent on the safe side in the VCA and increased descent time in the PDCT. Application of both the VCA and PDCT in mice can detect binocular dysfunction, which has high clinical relevance. However, the small effect sizes across VCA metrics limit the power and utility of this assay to detect partial effects. In contrast, the PDCT allows the use of a smaller sample size to detect binocular vision deficits, making it the assay of choice when studying stereoscopic depth perception in mice, and deficits that result from deprivation amblyopia.

